# Heroin Self-Administration Induces Altered Dopamine Terminal Dynamics in the Nucleus Accumbens

**DOI:** 10.1101/2022.02.02.478890

**Authors:** Brianna E. George, Monica H. Dawes, Emily G. Peck, Sara R. Jones

## Abstract

Administration of heroin results in the engagement of multiple brain regions and the rewarding and addictive effects are mediated, at least partially, through activation of the mesolimbic dopamine system. However, less is known about dopamine system function following chronic exposure to heroin. Withdrawal from chronic heroin exposure is likely to drive a state of low dopamine in the nucleus accumbens (NAc), as previously observed during withdrawal from other drug classes. Thus, we aimed to investigate alterations in NAc dopamine terminal function following chronic heroin self-administration to identify a mechanism for dopaminergic adaptations. Adult male Long Evans rats were trained to self-administer heroin (0.05 mg/kg/inf, IV) and then placed on a long access (FR1, 6-hrs, unlimited inf, 0.05 mg/kg/inf) protocol to induce escalation of intake. After one day of withdrawal, male rats exhibited lower basal extracellular levels of dopamine as well as reduced dopaminergic responses to a heroin challenge (0.1 mg/kg/inf, IV). Following heroin self-administration, rats had decreased basal extracellular levels of dopamine and blunted dopamine response following a heroin challenge (0.1 mg/kg/inf, IV) in the NAc compared to saline controls. FSCV revealed that heroin-exposed rats have reduced stimulated dopamine release during tonic-like, single-pulse stimulations but increased phasic-like dopamine release during multi-pulse stimulation trains (5 pulses, 5-100Hz) in addition to an altered dynamic range of release stimulation intensities when compared to controls. Further, we found that presynaptic D3 autoreceptor and kappa-opioid receptor activity was increased following heroin self-administration. These results reveal a marked low dopamine state following heroin-exposure and suggest the combination of altered dopamine release dynamics may contribute to increased heroin seeking.

## Introduction

The declaration of the present opioid overdose epidemic came as the result of the dramatic increasing prevalence of opioid use disorder (OUD) and opioid-related overdoses since the 1990s. In 2019, it was estimated that more than 130 people died each day from opioid overdose, with heroin involvement in nearly a third of those overdoses[1]. This overdose epidemic has been exacerbated by the social isolation and instability experienced during the COVID-19 pandemic, with preliminary reports from the CDC suggesting a 30% increase in opioid-related deaths in 2020[2]. Furthermore, disruptions in access to clinical providers and treatment as a result of COVID-19 restrictions have led to significant complications in opioid agonist treatment availability and distribution[3–5], highlighting the need for re-evaluation of current treatment options and expansion of preclinical research into the neurobiology of OUD to identify new treatment targets[4,6–9].

There are currently only three medications approved by the FDA to treat opioid use disorder: buprenorphine, methadone, and naltrexone. The most effective treatment for OUD is opioid replacement therapy, using either methadone or buprenorphine[10]. However, there are caveats with both medications: methadone has high abuse liability and can cause respiratory depression in patients[11] and buprenorphine is less successful in maintaining patient compliance when compared to methadone-assisted treatment[10]. Therefore, the need for novel, effective treatments with low abuse potential is substantial.

To find potential targets for OUD treatment, it is important to understand the mechanisms of action and consequences of chronic exposure to opioids. The prevailing hypothesis is that the reinforcing effects of exogenous opioids occur via disinhibition of dopamine neurons by activation of mu-opioid receptors (MOR) located on GABAergic neurons within the ventral tegmental area (VTA)[12–16]; however, some research has suggested that this model may not be the only mechanism involved in opioid reinforcement[17–21]. Support for the disinhibition hypothesis comes from numerous studies which show that opioid-induced disinhibition of VTA dopamine neurons leads to elevation of downstream dopamine release in the nucleus accumbens (NAc), which is a common mechanism of action across many drugs of abuse. Beyond initial reinforcing effects, chronic exposure to opioids is suggested to lead to persistent alterations in VTA dopamine neuron morphology and function. Chronic exposure to opioids, including morphine and heroin, has been shown to increase the basal firing rate and decrease the soma size of VTA dopamine neurons[22–27], while simultaneously decreasing extracellular dopamine levels[28] and levels of phosphorylated tyrosine hydroxylase[29] in the NAc. The range of hypotheses currently under investigation highlights the lack of understanding regarding neurophysiological alterations following chronic opioid exposure.

Thus, the present study aimed to identify functional and cellular changes following heroin self-administration in an attempt to establish a cellular mechanism through which chronic heroin exposure leads to reduced accumbal dopamine system function. Because hypodopaminergia promotes escalations in drug-seeking and propensity for drug relapse, pinpointing a mechanism for reduced dopamine function following heroin self-administration may lead to potential novel targets for treatments.

## Methods and Materials

### Animals

Adult male Long Evans rats (300-400g, Envigo, Indianapolis, IN) were given *ad libitum* access to food and water and maintained on a 12-hour modified reversed light/dark cycle (0300 hours lights off; 1500 hours lights on). Animal care procedures were in accordance with the National Institutes of Health guidelines in the Association for Assessment and Accreditation of Laboratory Animal Care and the experimental protocol was approved by the Institutional Animal Care and Use Committee at Wake Forest School of Medicine.

### IV Catheter Surgery

Rats were anesthetized with ketamine (100 mg/kg, i.p.) and xylazine (10 mg/kg, i.p.) and implanted with a chronic indwelling jugular catheter as described previously[30,31]. Immediately following the surgery, rats were administered meloxicam (1 mg/kg, s.c.) as a postsurgical anesthetic and single housed in custom chambers that functioned as both a housing cage and operant chamber.

### Heroin Self-Administration Procedures

Self-administration sessions were conducted during the active/dark cycle, between 0900 and 1500 hours. After at least two days of recovery, rats were trained to lever press for heroin on a fixed ratio one (FR1) schedule of reinforcement using a single active lever which, when pressed, delivered an intravenous infusion of heroin (0.05 mg/kg/inf). Acquisition self-administration sessions were initiated with the extension of the active lever into the chamber, with no experimenter-delivered priming infusions, and were terminated after 20 infusions or six hours, whichever occurred first. During the sessions, each lever response resulted in a 20-s time-out period in which the cue light was illuminated. Acquisition criteria were set as two consecutive daily sessions during which 20 infusions were obtained. When acquisition criteria were met, rats were placed on a long access paradigm with unlimited access to heroin (0.05 mg/kg/inf) on an FR1 schedule of reinforcement during 6-hour sessions occurring across 10 consecutive days.

### Ex Vivo Fast-Scan Cyclic Voltammetry

Fast-scan cyclic voltammetry (FSCV) was used to examine presynaptic NAc dopamine terminal release kinetics and dopamine autoreceptor and kappa opioid receptor (KOR) activity. Approximately 18 hours following the last long access session, rats were deeply anesthetized with isoflurane and rapidly decapitated. The brain was removed and immersed in oxygenated artificial cerebrospinal fluid (aCSF) containing (in mM): NaCl (126), KCl (2.5), NaH_2_PO_4_ (1.2), CaCl_2_ (2.4), MgCl_2_ (1.2), NaHCO_3_ (25), glucose (11), L-ascorbic acid (0.4). A vibrating tissue slicer (Leica VT1200S, Leica Biosystems, Wetzler, Germany) was used to prepare 400 μm thick coronal brain slices containing the NAc core. Slices were transferred to a recording chamber and submerged in a bath of oxygenated aCSF (32 °C) perfused at a rate of 1 mL/min. A carbon-fiber microelectrode (CFE; 100–200◻μM length, 7◻μM radius) and bipolar stimulating electrode were placed in the NAc core. Endogenous dopamine release was electrically evoked by a single pulse (750 μA, 4 msec, monophasic) applied to the slice every 3 minutes. Extracellular dopamine was measured by applying a triangular waveform (−0.4 to +1.2 to −0.4◻V *vs* Ag/AgCl, 400◻V/s) to the CFE. After an hour of slice acclimation and when the dopamine signal was stable for at least three successive collections, slices were used for one of the following experiments listed below.

### D2/D3 Autoreceptor and KOR Activity

> To probe activity of presynaptic autoreceptors in regulation of terminal release, cumulative concentrations of the D2 agonist, sumanirole (0.01, 0.03, 0.1, 0.3, 1, 3 μM), and the D3 autoreceptor and postsynaptic receptor (D3R) agonist, PD-128907 (0.001, 0.003, 0.01, 0.03, 0.1, 0.3 μM), were bath applied to separate slices. KOR activity was measured using the KOR agonist, U50-488 (0.01, 0.03, 0.1, 0.3, 1, 3 μM). Each slice was exposed to only one drug. The interstimulus interval for all concentration curves was three minutes.

### DA Release across Varying Stimulation Parameters

> The effects of two different stimulation patterns on dopamine release were tested. A frequency-response curve was obtained using 5 pulse stimulations at 5, 10, 20, 40, 60, and 100 Hz frequencies with 5-minute interstimulus intervals. A stimulation intensity curve was collected by using a single pulse stimulation at varying intensities, 50, 100, 150, 200, 250, 300, 350, 400, 450, 500, 600, 700, 800, 900, 1000 μA with an interstimulus interval of 60 seconds.

### FSCV Data Analysis

Demon Voltammetry and Analysis software[32] was used to analyze all FSCV data. Recording electrodes were calibrated at the end of every experiment by washing a known concentration of dopamine over the CFE using a flow-injection system and measuring the resulting current. The calibration factor of each electrode was used to convert the electrical current measured during experiments to dopamine concentration. Michaelis-Menten modeling was used to determine the concentration of dopamine released and the maximal rate of uptake (V_max_) following electrical stimulation.

### Microdialysis

Microdialysis surgeries were performed approximately 18 hours following the final long access self-administration session (Heroin n=5; Naïve n=5). Guide cannula (MD-2250; BASi Instruments, West Lafayette, IN) were stereotaxically implanted 2mm above the NAc (AP: +1.6mm; L: ± 1.4mm; DV: −6.0mm), as the probe extends 2mm past the cannula. Concentric microdialysis probes (2mm membrane, MD-2200; BASi Instruments, West Lafayette, IN) were inserted approximately 36 hrs post-surgery and continuously perfused with artificial cerebrospinal fluid (aCSF; pH 7.4; NaCl 148 mM, KCl 2.7 mM, CaCl_2_ 1.2 mM, MgCl_2_ 0.85 mM). For basal extracellular dopamine level experiments, overnight samples were collected at 0.5 μL/min from 1700 hours to 0800 hours, at which time the flow rate was increased to 1.0 μL/min. Baseline samples were collected every 20 minutes for two hours before intravenous administration of 0.1 mg/kg heroin. Following administration of heroin, 20-minute duration samples were collected for another two hours. All dialysate samples were frozen at −80° before determination of dopamine concentrations by high-performance liquid chromatography (HPLC) with electrochemical detection (ESA/Thermo Scientific, Chelmsford, MA)

### High-Performance Liquid Chromatography

All dialysate samples were analyzed using high-performance liquid chromography (HPLC) (ESA/Thermo Scientific, Chelmsford, MA) coupled with electrochemical detection at +220 mV using a high sensitivity analytical cell 5011A (Fisher) on a Choulechem III Electrochemical Detector (ESA Inc.). Neurotransmitters and their metabolites were separated on a Luna 100 × 3.0 mm C18 3 μm HPLC reversed-phase column (Phenomenex). The mobile phase consisted of 75 mM NaH2PO4, 1.7 mM 1-octanesulfonic acid sodium salt, 100 μL/L triethylamine, 25 μM EDTA, 10% acetonitrile v/v, at pH=3.0. Analytes were quantified using Chromeleon software (ESA/Thermo Scientific, Chelmsford, MA) by standards with known dopamine concentrations.

### Statistics

GraphPad Prism 7 (La Jolla, CA, USA) was used to statistically analyze all datasets and create graphs. Behavioral data were analyzed using either a Student’s t-test or two-way repeated-measures ANOVA followed by Bonferroni’s posthoc test where noted. Baseline voltammetry release and reuptake data, before and after drug perfusion, were compared using a one-way ANOVA, with group differences being tested using Tukey’s posthoc tests where noted. Phasic/tonic ratios and all western blot hybridization data were analyzed using a two-tailed Student’s *t*-test. All data are reported as mean ± SEM. All *p*-values of <0.05 were considered to be statistically significant.

## Results

### Long access heroin self-administration leads to escalation of heroin intake

During acquisition, rats reached acquisition criteria on average in 10 ± 1.072 days (mean ± SEM) (Fig. 1B). After acquisition criteria were met, rats were tested for heroin responding (0.05 mg/kg/inf) on an FR1 schedule of reinforcement during six-hour sessions with no limit on rewarded responses. Heroin was self-administered significantly more than saline (Fig. 1C, *F*(1,46) = 39.10, p<0.0001). Responding for heroin within the first hour increased across sessions (Fig. 1D, linear regression: β = 0.2483 ± 0.09339, *p* = 0.0083), and time to acquisition was negatively correlated with average responding for heroin across the last three sessions (Fig. 1E, β = −007987 ± 0.3890, *p* = 0.0499).

**Figure 1.**
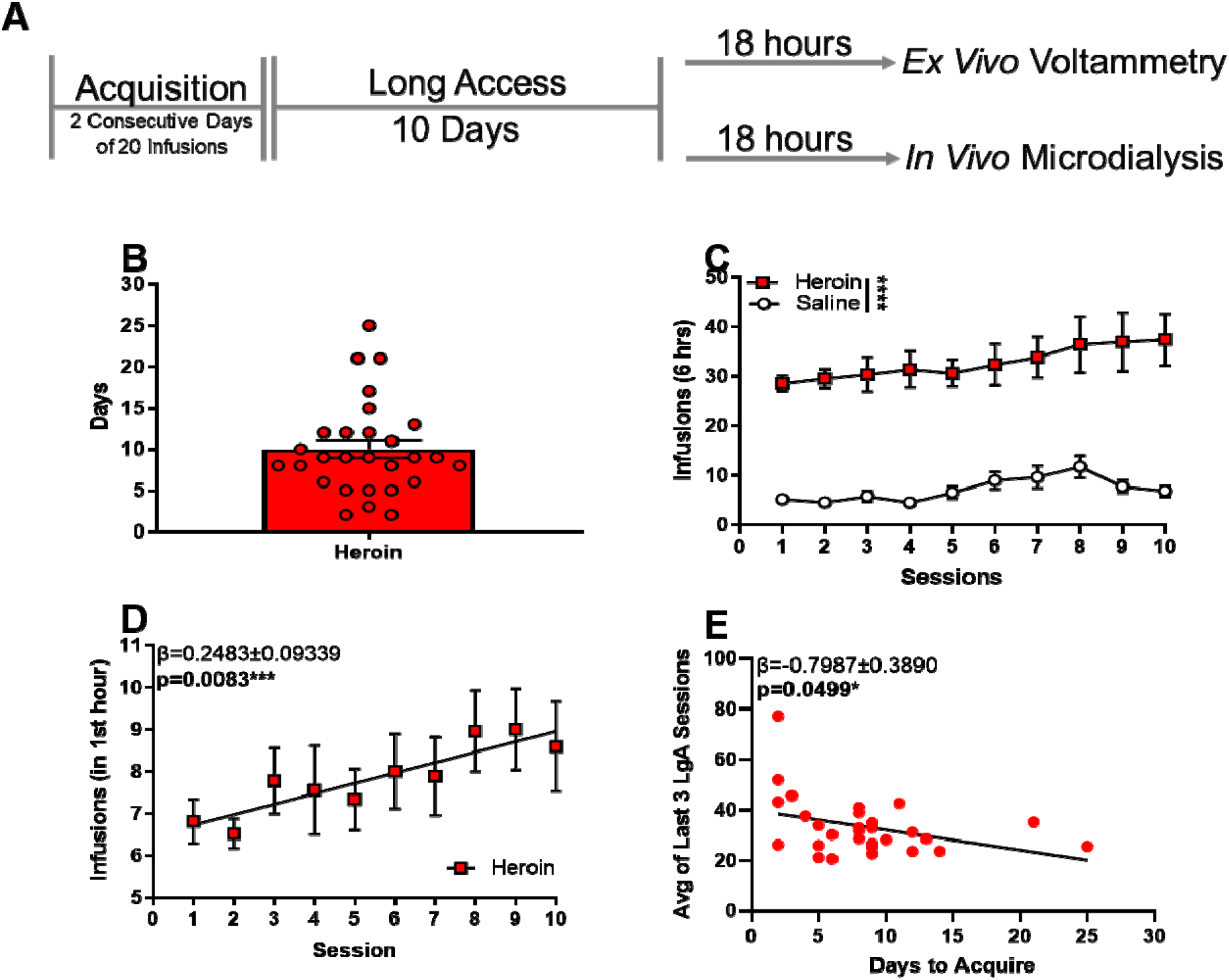
Long access heroin self-administration leads to escalation of heroin intake. (A) Experimental timeline of behavioral paradigm. (B) Total number of days to reach acquisition criteria. (C) Heroin intake during 6-hour long access sessions. (D) Heroin intake during the first hour of each session. (E) Correlation between the number of days to acquisition and average responding across the last 3 days of heroin long access self-administration. *p<0.05; **p<0.01; ***p<0.001; ****p<0.0001.

### Heroin self-administration reduced single-pulse stimulated dopamine release, but not dopamine reuptake in the NAc core

Rats were sacrificed for FSCV approximately 18 hours following the last heroin self-administration session. We found that single-pulse stimulated dopamine release was significantly lower in heroin exposed animals (Fig. 2B, t_62_ = 3.193, p = 0.0022), but that dopamine uptake was not significantly different between heroin exposed animals and saline controls (Fig. 2C, p > 0.05).

**Figure 2.**
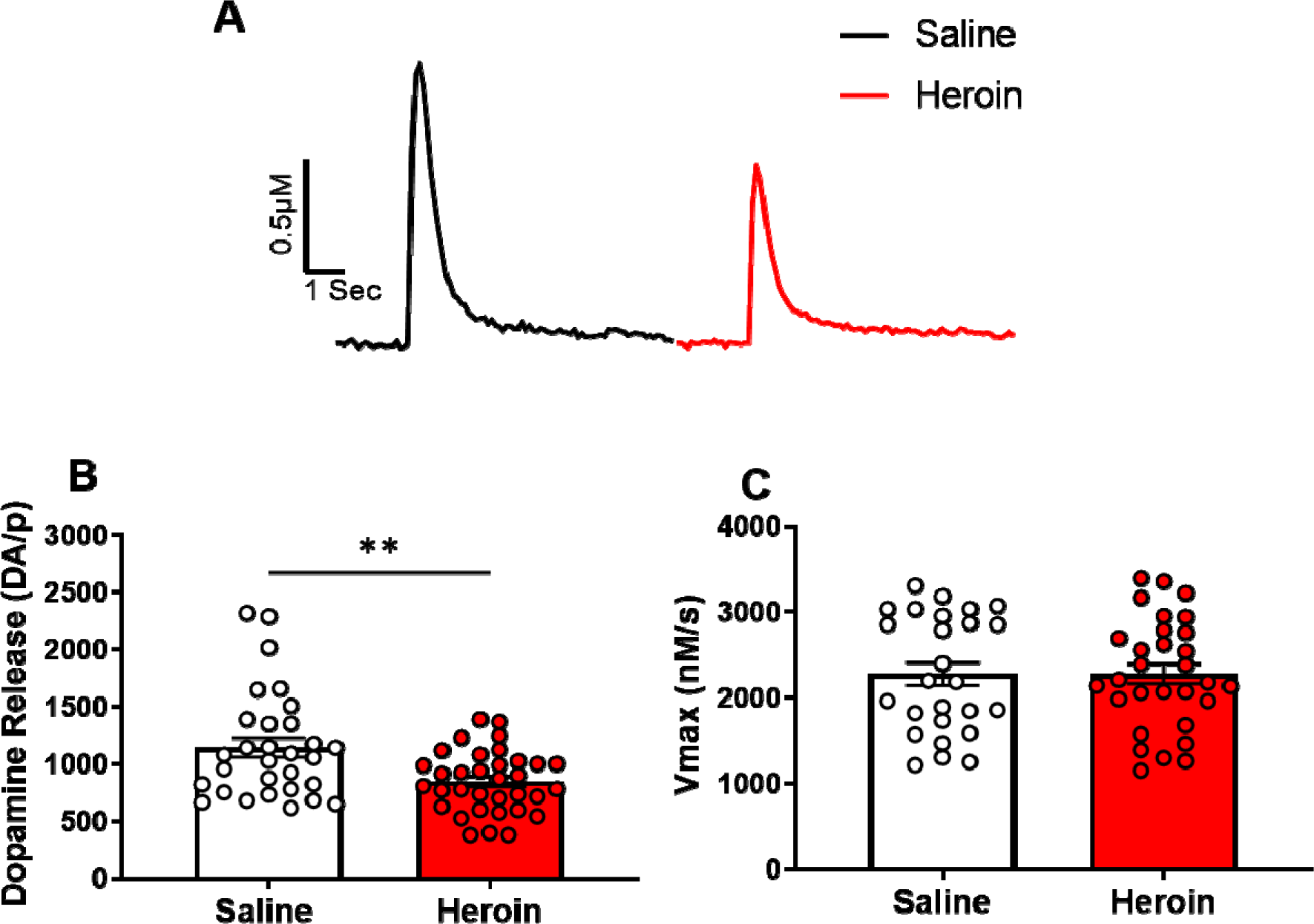
Heroin self-administration reduced single-pulse stimulated dopamine release, but not dopamine reuptake in the NAc core. (A) Representative traces of electrically-evoked dopamine in saline exposed (black) and heroin exposed (red) rats. (B) Stimulated dopamine release was significantly lower in heroin-exposed rats. (C) The maximal rate of dopamine uptake (Vmax) was not significantly different between saline and heroin exposed rats. *p<0.05; **p<0.01; ***p<0.001; ****p<0.0001.

### Opposite adaptations of D2 and D3 autoreceptors following heroin self-administration

To determine if heroin changed the activity of presynaptic D3 autoreceptors, regulation of dopamine release, we examined the release-inhibiting effects of the selective D3 receptor agonist, PD-128907. Two-way repeated-measures ANOVA revealed a significant main effect of heroin exposure (Fig. 3A, *F*(1,17) = 4.819, p=0.0423), such that heroin exposure resulted in decreased dopamine release, and increasing concentrations of PD-128907, which concentration-dependently inhibited dopamine release in both heroin and control animals (Fig. 3A, F(5,85) = 267.3, p<0.0001). Post-hoc analysis revealed a trend for greater inhibition in the heroin-exposed group at the 10nM (p=0.0556) and 30nM (p=0.0887) concentrations. In addition, the IC50 for PD-128907 was significantly decreased in heroin exposed rats (Fig. 3B, t_17_=2.317, p = 0.0332), indicating higher potency for inhibition; but the maximal effect at 300nM PD-128907 was not significantly different between heroin exposed and saline controls (Fig. 3C), suggesting no change in the overall efficacy of PD-128907.

**Figure 3.**
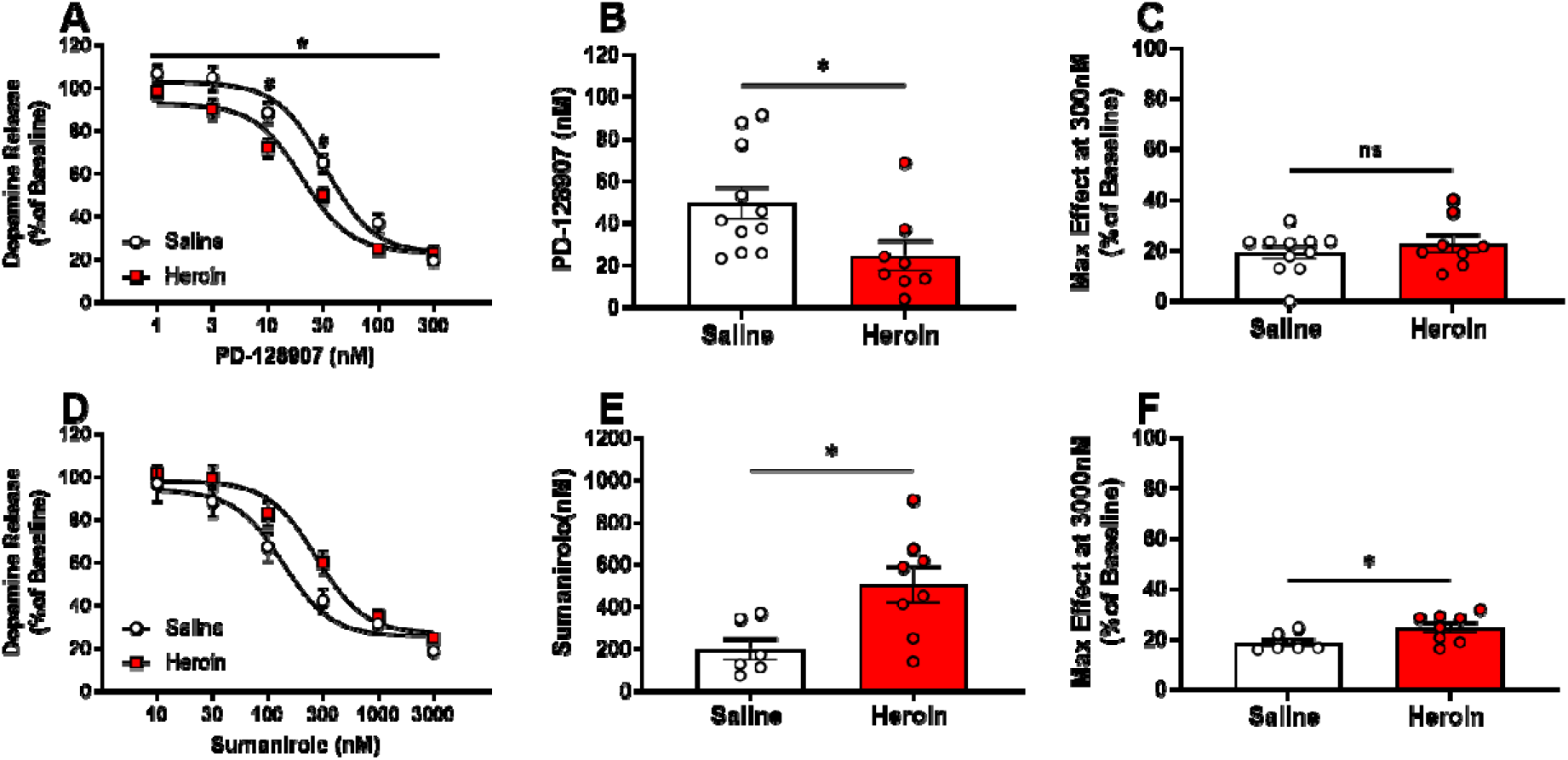
Opposite adaptations of D2 and D3 autoreceptors following heroin self-administration. Selective D3 agonist PD-128907 (A) decreased % baseline dopamine release in heroin exposed rats with (B) significantly lower IC50 in heroin exposed animals and (C) no significant differences in PD-128907 maximal effect (at 300 nM) between saline exposed and heroin exposed rats. Selective D2 agonist sumanirole (D) increased % baseline dopamine release in heroin exposed rats with (E) significantly greater IC50 in heroin exposed animals and (F) significantly greater maximal effect of sumanirole (at 3 μM) in heroin exposed rats. *p<0.05; **p<0.01; ***p<0.001; ****p<0.0001.

**Figure 4.**
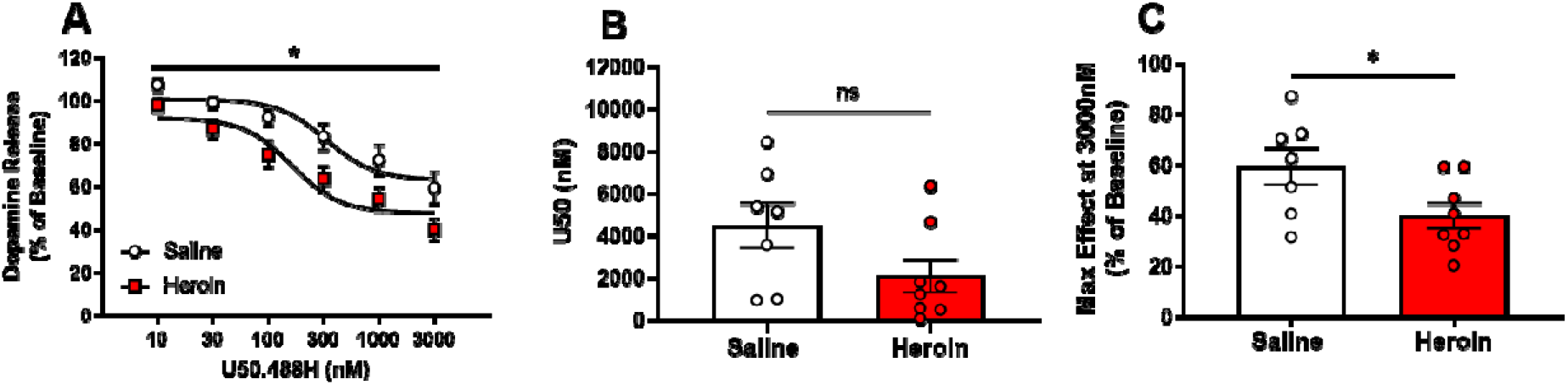
Presynaptic KOR activity on NAc dopamine terminals are increased by heroin self-administration. Selective kappa-opioid receptor agonist U50,488H (A) decreased % baseline dopamine release in heroin exposed rats with (B) no significant difference in IC50 between saline exposed and heroin exposed rats, but (C) significant decreases in the U50,488H maximal effect in exposed rats. *p<0.05; **p<0.01; ***p<0.001; ****p<0.0001.

To examine changes in D2 autoreceptor-mediated inhibition of dopamine release, the selective D2 receptor agonist, sumanirole, was used. Two-way repeated-measures ANOVA revealed a trend of the main effect of heroin exposure (Fig. 3D, F(1,12) = 3.213, p=0.0983), such that inhibition of stimulated dopamine by sumanirole may be reduced in heroin exposed rats. IC50 was significantly increased in heroin exposed rats (Fig. 3E, t_12_=2.774, p = 0.0168), as was the maximal effect of sumanirole (3 μM) treatment (Fig. 3F, t_12_=2.280, p = 0.0417).

### Presynaptic KOR activity on NAc dopamine terminals are increased by heroin self-administration

To determine the functional responsivity of KOR following heroin self-administration, the selective KOR agonist, U50,488H (U50), was used. U50 bath application resulted in a greater reduction of stimulated dopamine release in heroin-exposed rats than saline controls (Fig. 3G, F(1,13)=6.549, p=0.0238). Post-hoc analyses revealed a trend for decreased release in heroin exposed rats at the 300nM (p=0.0597), 1μM (p=0.0953), and 3μM (p=0.0632) concentrations. While the IC50 for U50 was not significantly different between heroin exposed rats and saline controls (Fig. 3H, p=0.0913), the maximal effect of 3μM U50 treatment was significantly decreased in heroin exposed rats (Fig. 3I, t_13_=2.226, p=0.0444).

### Heroin self-administration alters the dynamic range of dopamine release

MOR activation results in phasic dopamine firing, which is characterized by a “burst-like” pattern of increased action potentials at higher frequencies[33]. To examine how chronic heroin exposure alters dopamine terminal dynamics changes, we measured dopamine release under increased stimulation pulses (5 pulses) and increasing frequencies (5-100 Hz). When evaluating the absolute magnitude of dopamine release (μM) in response to multi-pulse stimulations, no significant group differences were observed (Fig 5A, p>0.05); however, heroin-exposed rats displayed a frequency-dependent increase in relative phasic release compared to naïve controls when dopamine release (μM) was normalized to single-pulse release (Fig 5B, F(1,20)=15.62, p=0.0008). Post-hoc analyses revealed significant differences at 40Hz (p=0.0009), 60Hz (p<0.0001), and 100Hz (p<0.0001). This effect was reflected by an augmented phasic/tonic ratio, which compares dopamine release evoked by 5 pulses at 60Hz with dopamine release evoked by a single pulse, in heroin-exposed rats (Fig 5C, t_23_=3.815, p=0.0009). These results suggest that chronic heroin self-administration may lead to reductions in basal dopamine release while enhancing the signal-to-noise ratio phasic release events, thereby causing dopamine dynamics to be phasic-shifted. This increase in the gain of dopamine terminal dynamics could lead to increased reinforcement of heroin self-administration and thus drive the increased escalation of intake.

**Figure 5.**
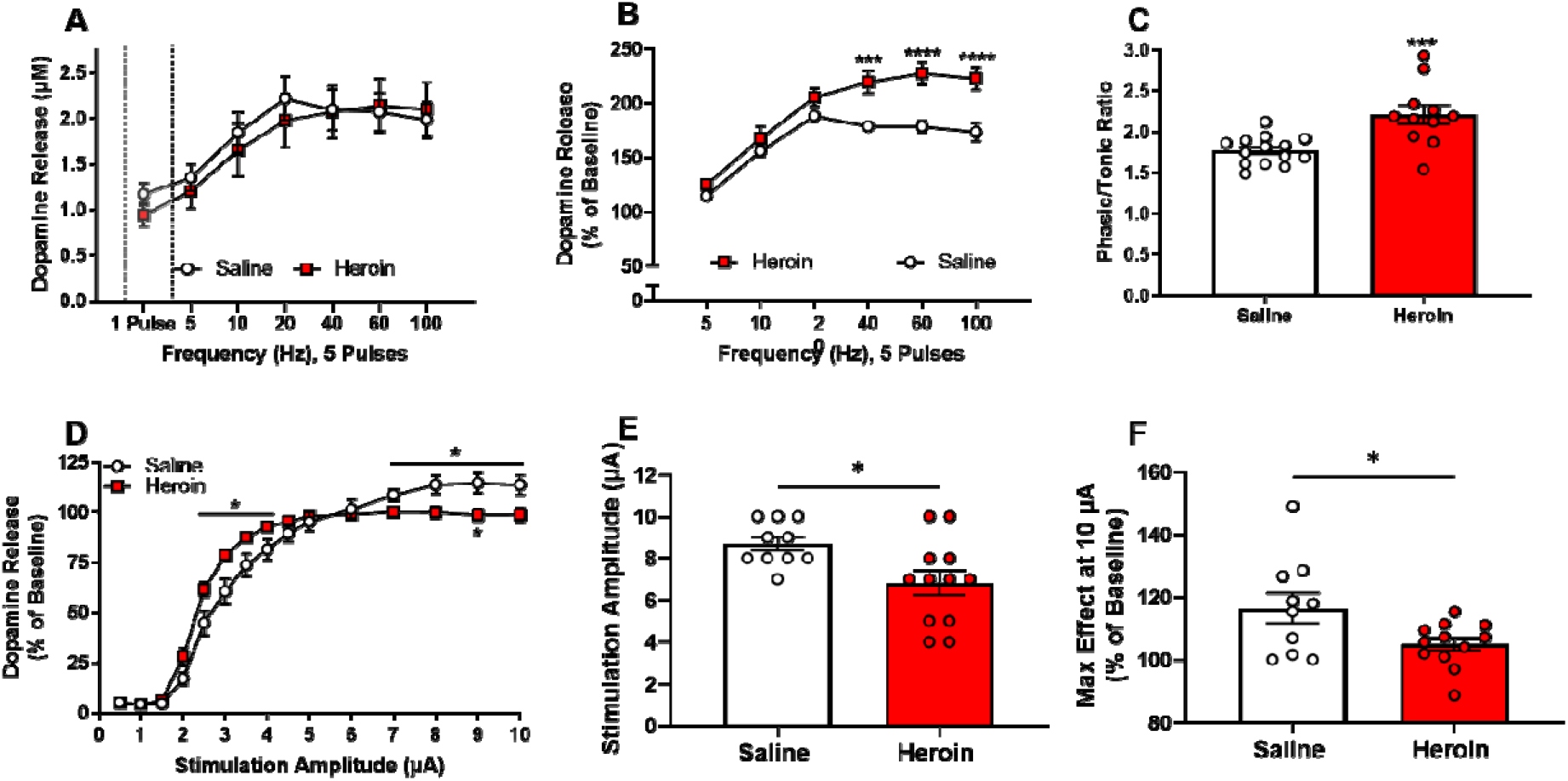
Heroin self-administration alters the dynamic range of dopamine release. Dopamine release was evoked with five pulses at increasing frequencies. (A) The magnitude of dopamine release did not differ between heroin-exposed and saline animals. (B) Percent dopamine release was significantly greater in heroin exposed rats at high frequencies. (C) The multi-pulse phasic/tonic ratio was significantly greater in heroin-exposed rats than saline exposed rats. (D) Dopamine release following single-pulse electrical stimulation of low and high intensities. Percent dopamine release was significantly greater at low (2.5 – 4 V) stimulation intensities and was significantly lower at high (7 – 10 V) stimulation intensities in heroin exposed rats. (E) The stimulation amplitude needed to elicit the greatest dopamine release was reduced in heroin-exposed rats. (F) The maximal release by the 10 μA stimulation was significantly decreased in heroin-exposed rats. *p<0.05; **p<0.01; ***p<0.001; ****p<0.0001.

**Figure 6.**
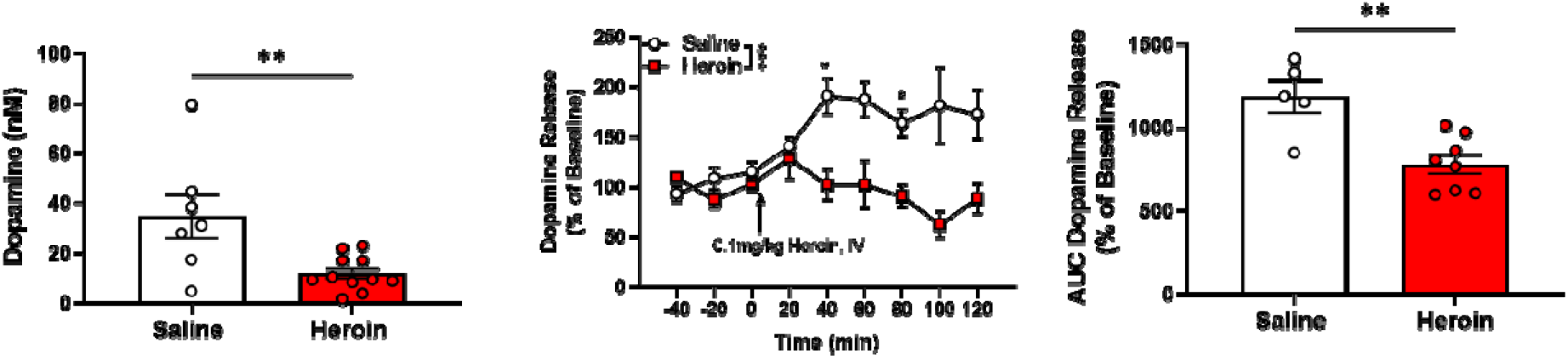
Heroin self-administration decreased basal extracellular levels of dopamine in the NAc and dopamine response to a heroin challenge. (A) Overnight (basal) extracellular dopamine was significantly lower in heroin-exposed rats. (B) Following the heroin challenge (0.1 mg/kg, IV), heroin exposed rats showed no significant increase in extracellular dopamine, while significant, prolonged increases were observed in saline rats. (C) Heroin exposed rats showed significantly decreased dopamine when compared to saline exposed rats. *p<0.05; **p<0.01; ***p<0.001; ****p<0.0001.

In addition to examining release under “phasic-like” stimulation parameters, we also measured dopamine release by altering the electrical stimulation amplitude. Heroin self-administration increased dopamine release elicited at low stimulation amplitudes and decrease of dopamine release elicited by high stimulation intensities (Fig. 5D). A two-way repeated-measures ANOVA of the low stimulation amplitudes (2.5-4μA) revealed significant differences between the heroin and saline groups (Fig. 5D, F(1,21)=5.494, p=0.0290). This effect was quantified by measuring the individual amplitude needed to elicit the greatest amount of dopamine for each slice, which showed that heroin self-administering animals required lower stimulation amplitudes to drive maximal dopamine release across stimulation intensities tested (Fig.5E, t_20_=2.611, p=0.0167). A two-way repeated-measures ANOVA of the high stimulation amplitudes (7-10μA) revealed significant differences between the heroin and saline groups (Fig. 5D, F(1,21)=7.519, p=0.0122) and post-hoc analyses showed a significant difference at the 9 μA stimulation amplitude (p=0.0442). When measuring the amount of dopamine released at the maximal stimulation amplitude (10 μA), we found heroin self-administering animals had lower dopamine release compared to saline controls (Fig.5F, t_20_=2.292, p=0.0329). Overall, these results suggest that the dynamic range of dopamine release across varying stimulation intensities is reduced following heroin self-administration. animals.

### Heroin self-administration decreased basal extracellular levels of dopamine in the NAc and dopamine response to a heroin challenge

Following one day of recovery after cannula implantation, microdialysis probes were inserted into the NAc, and aCSF was perfused at a low flow rate of 0.5 μL/min. A single 15-hour baseline sample was collected and analyzed for dopamine at equilibrium between the probe and extracellular tissue, to quantify basal extracellular levels. Approximately 24 to 32 hours following the last self-administration session, heroin-exposed animals had a significant decrease in basal levels of extracellular dopamine when compared to saline exposed controls (Fig. 5A, t_16_=3.052, p=0.0076). The next morning, the flow rate was increased to 1.0 μL/min to allow more rapid (20 min) sampling times. Following the collection of 6 total samples where dopamine levels were stable (<10% difference and no trend up or down) for at least 3 collections, all rats were administered 0.1 mg/kg heroin (IV), and samples were collected every 20 minutes for an additional 120 minutes. Extracellular dopamine levels were significantly increased in control rats following heroin administration, but there was no change in heroin exposed rats (Fig. 5B, *F*(1,12)=18.79, p=0.0010), with post-hoc analyses revealing significant differences at the 40 (p=0.0293) and 80 (p=0.0128) minute time points. Analysis of the area under the curve for the magnitude of dopamine outflow showed dopamine release was significantly lower in heroin exposed animals than in saline controls (Fig. 5C, t_11_=3.840, p=0.0027).

## Discussion

The current study demonstrated that a history of chronic long access heroin self-administration in rats induced adaptations in extracellular levels, terminal release dynamics, and presynaptic regulators of the dopamine system that are with an overall state of hypodopaminergia. Low dopamine activity following chronic substance use has been documented with alcohol, cocaine, amphetamine, and other stimulants in multiple species, including non-human primates and humans, but the impact of opioids such as heroin is less well-documented. Understanding the cellular mechanisms underlying heroin-induced hypodopaminergia will be important in combating OUD since low dopamine function is strongly associated with increased propensity to seek and take drugs as well as increased probability of relapse after abstinence.

When given access to heroin under long access parameters, male rats escalate their intake across sessions, as shown previously[34]. The literature shows that rodent heroin self-administration protocols lead to tolerance, withdrawal symptoms, and drug-induced symptomology similar to that observed in clinical populations with OUD. Therefore, we suggest that heroin self-administration is a useful model to evaluate heroin-induced alterations in neurobiology.

Following heroin self-administration, rats display reduced basal extracellular levels of dopamine and reduced dopamine release in response to single-pulse electrical stimulations, which are considered classical signs of reduced dopamine system function, or hypodopaminergia[35]. The present findings are in agreement with previous studies, showing that chronic exposure to opioids leads to reduced extracellular levels of dopamine in the striatum[36–38]. Furthermore, we found that an acute IV heroin challenge, which normally increases extracellular levels of dopamine, had no effect on dopamine in self-administering animals. This tolerance is consistent with self-reports from OUD sufferers who report that, after months or years of daily use, heroin no longer makes them “high”. Combined, these results suggest that chronic heroin exposure, like exposure to other drugs of abuse, induces a hypodopaminergic state. Hypodopaminergia has been linked to lead to a compensatory increase in drug-seeking behaviors and negative affect during drug withdrawal[39]. Thus, it is possible that the low functioning dopamine system observed following heroin self-administration exposure may contribute to the escalation of heroin seeking seen during long access.

Here we report that, after extended heroin self-administration, D2 autoreceptors show a decrease in function and/or activity, while D3 autoreceptors show increased function. D2-type autoreceptors modulate the firing rate of dopamine neurons, dopamine release, and synthesis of dopamine and are expressed on the soma, dendrites, and axon terminals of dopamine-rich regions such as the NAc and VTA. While a large body of literature exists supporting the role of D2 autoreceptors, evidence for the functionality of D3 receptors is contradictory[40,41]. Due to structural similarities between D2 and D3 receptors and the 10-fold greater expression of D2 receptors, the variability in support D3Rs as autoreceptors may be due to technical limitations. While acknowledging that most D2R and D3R selective agonists, including the ones used here, sumanirole and PD-128907, will act on both receptors at high concentrations, there were shifts in the potency of both drugs after heroin self-administration, but in opposite directions, suggesting a functional dichotomy between the two receptors. Similarly, other studies have reported opposing D2R and D3R changes in mesolimbic regions following exposure to opioids in rodents[42–45] and humans[46,47]. The results of this study are intriguing in light of other studies which show that D3 receptors antagonists are effective at reducing opioid self-administration[48–51] (for review see [52,53]). Indeed, it seems likely that D3R antagonists may work at both presynaptic and postsynaptic receptors to induce increased basal dopamine tone and block synaptic dopamine transmission, respectively, thus combining to reduce opioid seeking[54]. To our knowledge, this is the first report showing evidence of a functional increase in D3 autoreceptors in the NAc after opioid self-administration.

Similar to D3 autoreceptor activity, KOR activity was increased following heroin self-administration. KORs are located in the NAc on presynaptic dopamine terminals and, upon activation with the endogenous ligand (dynorphin), strongly inhibit dopamine release[55]. Prior studies have shown that dynorphin and its precursors are elevated following opioid exposure[56–60], which suggests increased endogenous agonist activation of KORs. Previous research has shown that administration of the KOR antagonist, norbinaltorphimine (nor-BNI), is effective at reducing heroin self-administration and withdrawal-induced negative affect[61]. Furthermore, another study reported that co-administration of heroin and the KOR agonist U50 reduced heroin-induced NAc dopamine release during self-administration and low doses of U50 increased heroin self-administration. Combined, this data suggests that inhibition of dopamine release through endogenous activation of the KOR as a result of chronic exposure may decrease the reinforcing properties of heroin and drive increased opioid seeking, particularly during withdrawal. The results of the current study expand upon previous findings, showing that KOR activity is increased after chronic opioid administration, and providing further evidence that KOR antagonist may be a viable treatment option for OUD.

Perhaps the most surprising discovery was that following heroin self-administration, dopamine release in the NAc under “phasic-like” electrical stimulation was increased. However, when we elicited single-pulse dopamine release at increasing stimulation intensities, heroin-exposed rats had greater release than the control group at low amplitude stimulation, but reduced release at high amplitude stimulation. A previous study has shown that MOR activation results in presynaptic facilitation in the NAc; likely through inhibition of cholinergic interneuron activity as seen with nicotinic antagonists[62]. This presynaptic facilitation results in reduced dopamine release under single-pulse stimulations and subsequent enhancement of dopamine release under burst-like stimulations due to increased accumulation of presynaptic calcium. Thus, it is possible that the mechanisms by which nicotine and opioids regulate dopamine release in the NAc have some functional overlap[62]. This idea is further supported by evidence showing that chronic nicotine exposure leads to reduced single-pulse dopamine release, increased paired-pulse ratios, and increased phasic/tonic ratios in the NAc[63,64]. This is consistent with our findings as, when normalized to single-pulse release, we observed increased dopamine release under multi-pulse stimulations in heroin-exposed animals compared to saline controls; however, the absolute magnitude of dopamine release under single-pulse stimulations was reduced in heroin-exposed animals compared to saline counterparts. This finding is enhanced by results from the stimulation intensity experiment, which shows that dopamine release was facilitated at low stimulation intensities and decreased during high stimulation intensities during single-pulse stimulations. While this finding may seem counterintuitive, this may suggest that the dynamic range of dopamine release is largely reduced following chronic heroin exposure. In a similar manner to the previous experiment, while lower stimulation intensities increased release when normalized to their single-pulse baseline release in heroin-exposed animals because their basal levels of dopamine are reduced, the overall magnitude of dopamine release is still reduced in heroin-exposed animals compared to naïve counterparts. This concept coupled with the finding that dopamine release is reduced under high stimulation intensities in heroin rats suggests a restricted range of activity-dependent dopamine release which has been previously shown to occur following chronic nicotine exposure and is thought to contribute to altered coding of dopamine reward signaling[65].

Our group has recently shown that extended heroin self-administration leads to increased dopamine release under ‘burst-like’ stimulation in female, but not male, rats[30]. While these previous results may appear contradictory when compared to current findings, some methodological differences may account for differing results. In the previous study, dopamine recordings were conducted in the medial NAc shell while the current study recorded in the NAc core. The NAc shell is known to have different dopaminergic innervation[66,67] and distribution of dopamine regulators such as autoreceptors, KOR, and DAT than the NAc core[68], suggesting that variances in the regulation of dopamine transmission may account for some of the observed differences[69]. Second, in the prior study, all rats self-administered a lower heroin dose than in the current study (0.025 mg/kg/inf vs. the current 0.05 mg/kg/inf). The upward shift in heroin dose-responsiveness in female rats is observed across various doses (indicating increased efficacy); therefore it is possible that, in males, the lower dose used previously was not sufficient to elicit the adaptions to phasic dopamine release seen in this experiment. While the current study did not utilize females, given previous findings, it is possible that female rats may exhibit even greater dopaminergic adaptations than what is seen in males here.

Overall, this study builds support for a mechanism in which chronic exposure to heroin results in reduced dopamine system function and highlights potential targets for pharmacological interventions, presynaptic D3 receptors, and KOR. Therapeutics aimed at restoring dopamine levels after chronic opioid exposure may be beneficial in treating OUD. While the current study measured outcomes during what would commonly be considered an “early withdrawal” time point, it would be interesting to evaluate if these alterations in dopamine function persist during late withdrawal.

